## Supplementary material for "Lytic transglycosylases mitigate periplasmic crowding by degrading soluble cell wall turnover products": Fig1 Source Data 1: UncroppedWesterns.pdf

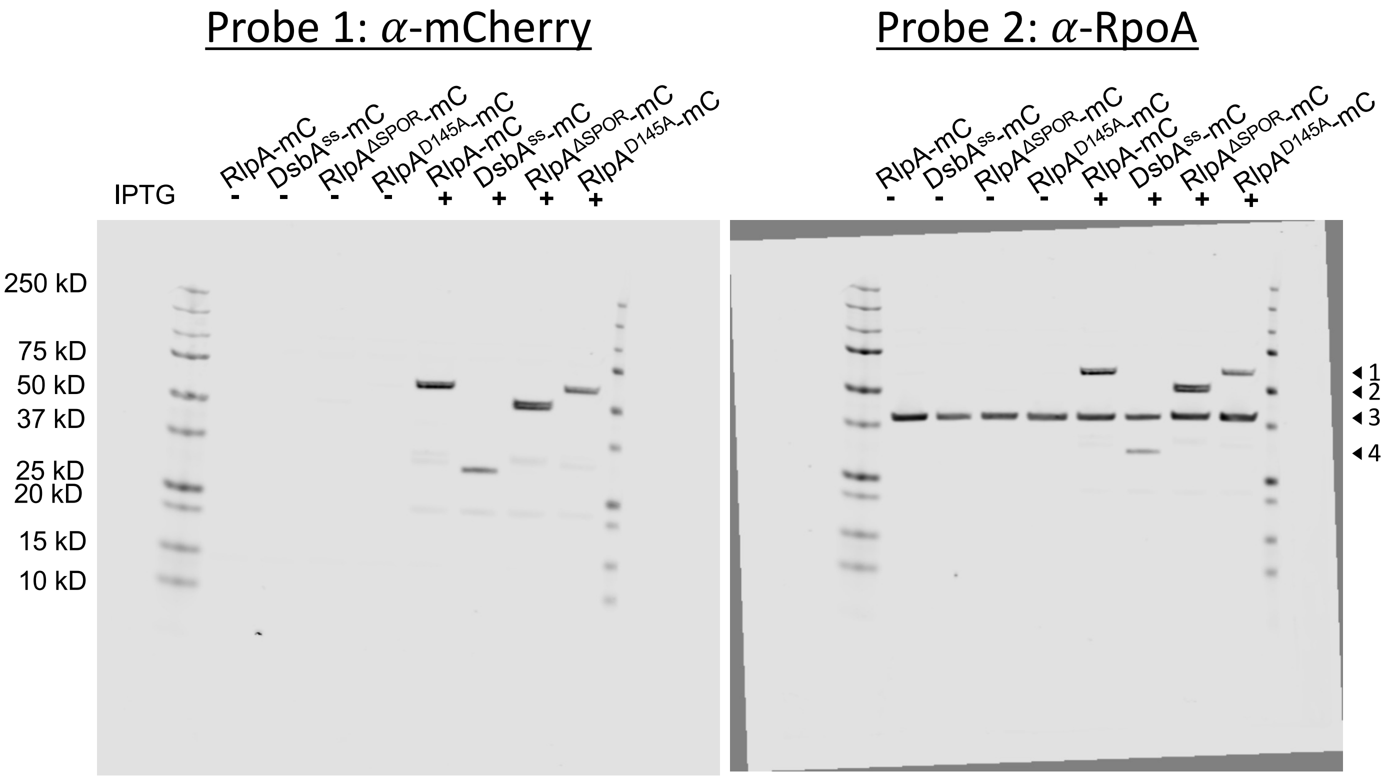


| Major Band | Annotation | Approx. Predicted Size |
|---|---|---|
| 1 | Full length RlpA-mCherry fusion | 55 kD |
| 2 | $RlpA^{\Delta SPOR}$-mCherry fusion | 46 kD |
| 3 | RpoA | 37 kD |
| 4 | $DsbA^{ss}$- mCherry<br>(Mature, secreted and signal peptide-cleaved) | 27 kD |
